# Spiral brain dynamics track the temporal unfolding of subjective intensity during the *N,N*-dimethyltryptamine experience

**DOI:** 10.64898/2026.09.22.753116

**Authors:** Gabriela Sawicka, Marian Martínez-Marín, Christopher Timmermann, Robin Carhart-Harris, Morten Kringelbach, Yonatan Sanz-Perl, Jakub Vohryzek, Gustavo Deco

**Author notes:** These authors contributed equally (Gabriela Sawicka, Marian Martínez-Marín). These authors jointly supervised this work (Jakub Vohryzek, Gustavo Deco).

## Abstract

Psychedelic compounds such as N,N-dimethyltryptamine (DMT) induce rapid and profound changes in conscious experience, requiring neuroimaging measures that capture how brain dynamics evolve across space and time. Emerging evidence suggests that rotational (vortex-like) dynamics may contribute to the coordination of large-scale neural activity. Here, we introduce two vorticity-based metrics; spiral vorticity (SV) and spatial spiral vorticity dispersion (SSVD), to quantify the rotational organization and spatial heterogeneity of large-scale cortical phase dynamics. Using fMRI data acquired under placebo and DMT in a within-subject design, we assessed these measures globally, across resting-state networks, and regionally to examine their relationship to subjective intensity.

DMT increased SV and SSVD and reduced temporal irreversibility, indicating stronger rotational organization, greater spatial heterogeneity, and altered directional structure of cortical dynamics. Effects were non-uniform across functional systems, with the largest changes in default mode, visual, frontoparietal, and dorsal attention networks. Both vorticity measures tracked the temporal evolution of subjective intensity globally, while network-level analyses revealed spatially heterogeneous brain–experience coupling. Spatiotemporally resolved analyses further showed that the strongest reorganization of spiral dynamics emerged during offset and recovery rather than at peak subjective intensity.

These findings identify spiral dynamics as a time-resolved neural signature of the DMT experience. By quantifying the rotational structure and spatial heterogeneity of cortical phase flow, vorticity-based measures provide a framework for linking large-scale brain dynamics to evolving subjective experience.

## Introduction

A central challenge in neuroscience is to explain how time-varying brain activity relates to the unfolding of subjective experience. Large-scale neuroimaging has therefore moved beyond static summaries toward time-resolved descriptions of neural dynamics (Vidaurre et al. 2017; Zalesky et al. 2014; Preti et al. 2017; Cabral et al. 2017), creating new opportunities to link evolving brain states to moment-to-moment changes in consciousness.

Classic serotonergic psychedelics provide a powerful model for studying this relationship because they induce marked, quantifiable changes in both brain activity and subjective experience. They have also attracted growing clinical interest (Garcia-Romeu et al. 2016). Intravenous *N,N*-dimethyltryptamine (DMT) is particularly well suited to time-resolved brain–experience analyses: its effects emerge rapidly, peak within approximately 2–3 minutes, and largely resolve within 15–30 minutes, depending on dose and administration protocol (Timmermann et al. 2019; Strassman 1995; Vogt et al. 2023). Here, we take advantage of this temporal profile to test whether evolving cortical dynamics track the unfolding intensity of the DMT experience.

DMT rapidly alters large-scale functional organisation. Functional magnetic resonance imaging (fMRI) studies show that DMT alters global connectivity, network segregation (Timmermann et al. 2023; Girn et al. 2026), cortical hierarchy (Vohryzek, Garcia-Guzmán, et al. 2025), and transmodal organisation (Vohryzek, Cabral, et al. 2024). Several of these changes track subjective intensity, as do electrophysiological measures of oscillatory activity, signal diversity, and travelling waves (Timmermann et al. 2019; Lewis-Healey et al. 2026; Alamia et al. 2020).

Together, these findings show that the temporal evolution of the DMT experience is reflected in large-scale brain dynamics. However, it remains unclear how cortical activity flow propagation and reorganisation shape this distinctive spatiotemporal brain activity landscape. Here, we address this gap by examining rotational phase-flow dynamics in whole-brain fMRI. Rotational and wave-like neural dynamics have been observed across spatial scales, from mesoscopic cortical recordings (Rubino et al. 2007; Townsend et al. 2015; Huang et al. 2010; Wu et al. 2008) to macroscopic whole-brain activity measured with EEG (Blackburne et al. 2025; Freeman and Barrie 2000), electrocorticography (ECoG) (Muller et al. 2016), MEG (Ribary et al. 1991), and fMRI (Xu et al. 2023). Such dynamics have been implicated in memory (Das et al. 2026), sleep (Muller et al. 2016), and cognitive processing, with emerging evidence suggesting that rotational organisation may play an important mechanistic role in coordinating large-scale brain activity (Xu et al. 2023; Ye et al. 2026).

In the present study, we test whether rotational organisation changes under DMT and tracks the temporal evolution of subjective intensity. Our approach builds on phase-based descriptions of large-scale brain activity. The global Kuramoto order parameter quantifies whole-brain phase synchronisation, while its temporal variability indexes global metastability (Cabral et al. 2014; Deco et al. 2017; Deco and Kringelbach 2020). Its local extension defines a spatially resolved synchronisation field whose continual reorganisation across space and time forms the basis of the turbulence framework (Deco and Kringelbach 2020; Escrichs et al. 2022). Local synchronisation, however, does not directly describe the direction of cortical phase flow.

To capture this feature of brain dynamics, we analyse spatial gradients of the cortical phase field (Townsend et al. 2015; Townsend 2018; Xu et al. 2023) and quantify their rotational component as spiral vorticity (SV). We further quantify the spatial dispersion of SV across cortex as spatial spiral vorticity dispersion (SSVD). These measures complement local synchronisation: spiral structures are organised around phase singularities with low local Kuramoto order, while vorticity captures the coherent rotational organisation surrounding those centres (Deco et al. 2026).

Turbulence-based measures describe how local synchronisation evolves, but not how cortical dynamics are directionally orchestrated. To address this complementary property, temporal irreversibility has been used to quantify departures from detailed balance and infer directed functional hierarchy. This framework has provided mechanistic insight into differences in brain dynamics reorganisation in depression between psilocybin and escitalopram (Deco et al. 2024) and acute hierarchy flattening under psilocybin (Pasquini et al. 2024), LSD, and DMT (Vohryzek, Kringelbach, et al. 2024). We extend this approach to cortical phase flow by defining temporal irreversibility as the asymmetry between forward and time-reversed phase-flow transitions, indexing local departures from temporally symmetric dynamics.

We test three hypotheses. First, we ask whether DMT alters large-scale brain dynamics indexed by SV, SSVD, and temporal irreversibility. Second, we test whether SV and SSVD track subjective intensity. Finally, we examine whether these effects are spatially and temporally specific and exhibit structured patterns of cortical reorganisation across the unfolding DMT experience.

## Materials and Methods

### DMT dataset

Full details of participant characteristics, experimental design, and data acquisition are reported by (Timmermann et al., 2023). All participants provided written informed consent prior to participation. The study received approval from the National Research Ethics Committee London-Brent and the Health Research Authority and was conducted in accordance with the revised Declaration of Helsinki (2000), International Council for Harmonisation Good Clinical Practice guidelines, and the National Health Service Research Governance Framework. The study was sponsored by Imperial College London and conducted under a Home Office licence for Schedule 1 drugs.

### Participants

Twenty-five participants were recruited into a single-blind, counterbalanced, placebo-controlled study. Participants underwent physical and mental health screening, including physical examination, electrocardiography, blood pressure and pulse measurements, blood tests, and psychiatric assessment. Exclusion criteria included age below 18 years, no previous psychedelic or hallucinogen use, personal psychiatric history, immediate family history of psychosis, excessive alcohol consumption (> 40 units per week), and blood or needle phobia. Urine drug screening and, where applicable, pregnancy testing were also performed.

Twenty participants completed the study (7 female; mean age = 33.5 years, SD = 7.9). Three were excluded because of excessive motion during the 8-minute DMT scan (> 15% of volumes scrubbed at a framewise displacement (FD) threshold of 0.4 mm), leaving 17 participants, consistent with previous analyses of this dataset (Vohryzek, Cabral, et al. 2024). For the time-resolved analyses in Figures 3 and 4, three additional participants were excluded because > 20% of volumes were scrubbed using an FD threshold of 0.476 mm.

**Figure 1.**
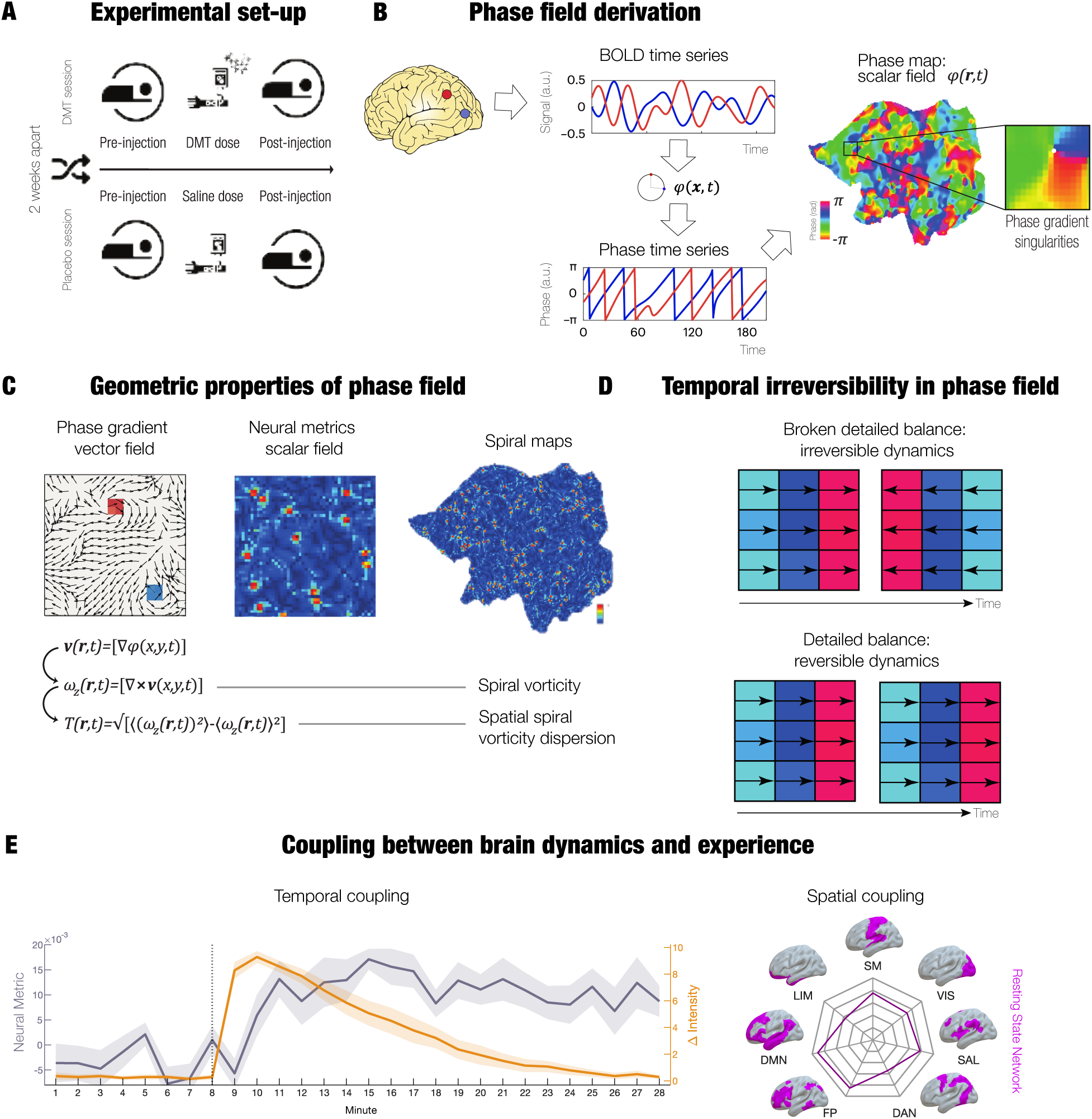
Methodological overview. **(A) Experimental setup.** Participants completed placebo and DMT fMRI sessions in a randomized crossover design separated by two weeks. During each session, intravenous saline or DMT was administered after an 8-minute baseline period. **(B) Phase field derivation.** BOLD fMRI signals were projected from 2-mm MNI volumetric space to HCP surface-vertex space and interpolated onto a regular two-dimensional cortical flatmap. The instantaneous phase of each grid element was estimated using the Hilbert transform, yielding a time-resolved cortical phase field, *φ*(***r***, *t*). Insets illustrate the local phase structure surrounding a phase singularity. **(C) Geometric properties of the phase field.** Spatial gradients of the phase field were used to construct a normalized phase-gradient vector field (left), ***v***(***r***, *t*) = *▽φ*(***r***, *t*). The local rotational component of this field was quantified by its curl, *ω_z_*(***r***, *t*) = [*▽* × ***v***(***r***, *t*)]*_z_*, defined here as spiral vorticity (SV). The spatial dispersion of vorticity values across cortical locations was quantified as spatial spiral vorticity dispersion (SSVD). Representative scalar-field maps (middle) and cortical renderings (right) illustrate these measures. **(D) Temporal irreversibility.** Temporal irreversibility was estimated from directional asymmetries between forward and reverse phase-flow transitions. Breaking detailed balance produces unequal forward and reverse transition probabilities, indicating irreversible dynamics, whereas balanced transitions indicate greater temporal reversibility. **(E) Coupling between brain dynamics and experience.** DMT-induced changes in spiral dynamics were related to subjective intensity using complementary analyses. Temporal coupling was assessed by correlating minute-wise changes in neural metrics with changes in subjective intensity, whereas spatial coupling was assessed using resting-state-network linear mixed-effects models relating within-subject fluctuations in neural dynamics to subjective intensity.

**Figure 2.**
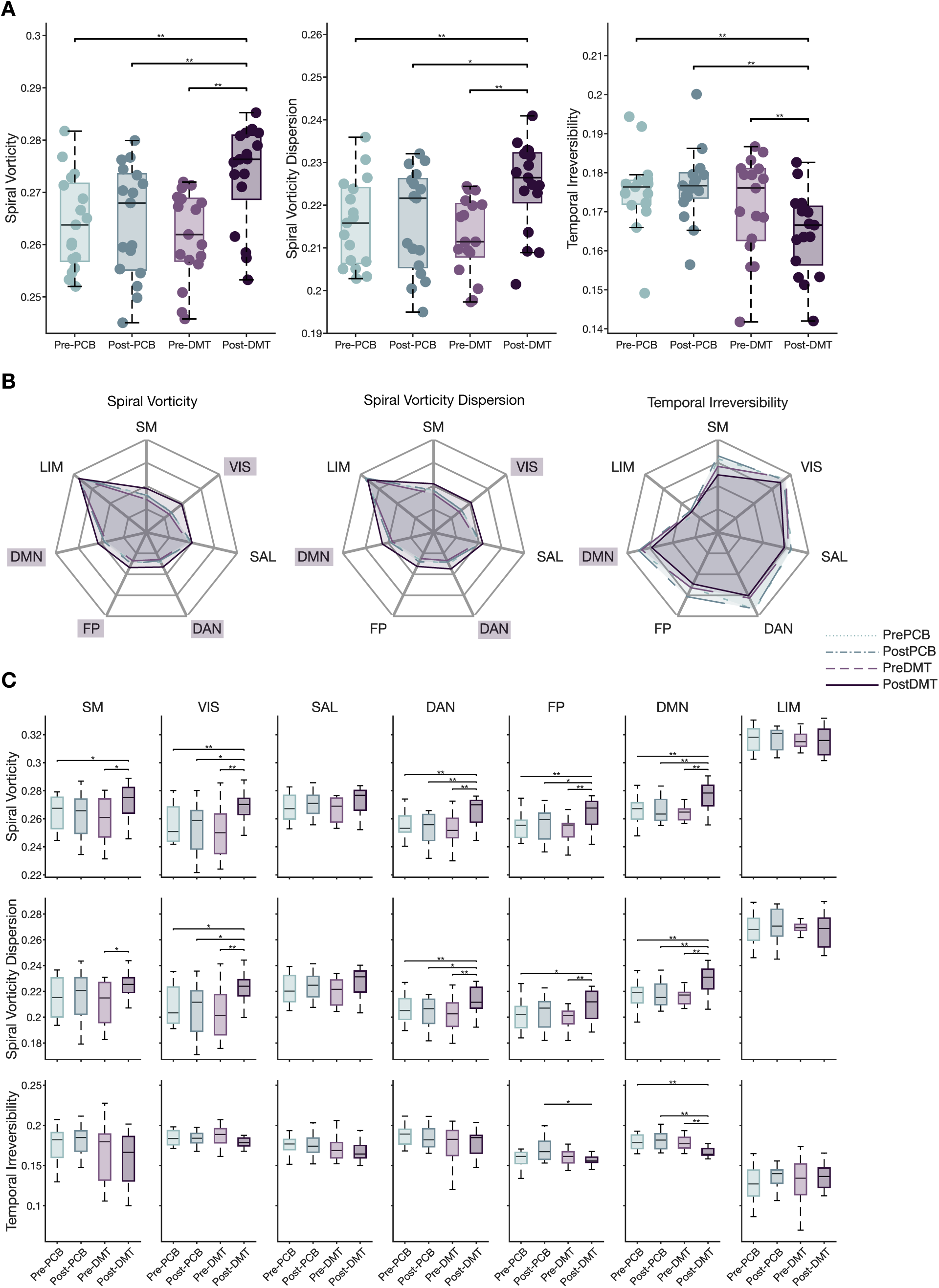
DMT alters global and network-specific brain dynamics. **(A) Global effects.** Group distributions of three neural dynamics metrics (left to right: Spiral Vorticity, Spiral Vorticity Dispersion, and Temporal Irreversibility) across four experimental conditions (Pre-PCB, Post-PCB, Pre-DMT, and Post-DMT). **(B) Network-level summaries.** Radar plots display the median value of each metric within the Yeo 7-network parcellation for each condition. Network labels are highlighted in purple where the Post-DMT condition differs significantly from all other conditions (Pre-PCB, Post-PCB, and Pre-DMT). **(C) Network-level statistical comparisons.** Boxplots show condition-wise distributions for each resting-state network and significant pairwise differences (* *p_FDR_* < 0.05, ** *p_FDR_* < 0.01, *** *p_FDR_* < 0.001; paired sign-flip permutation test). Global analyses were Benjamini–Hochberg FDR-corrected across the six pairwise condition comparisons within each metric. Resting-state-network analyses were FDR-corrected across six pairwise condition comparisons and seven networks (42 comparisons per metric).

**Figure 3.**
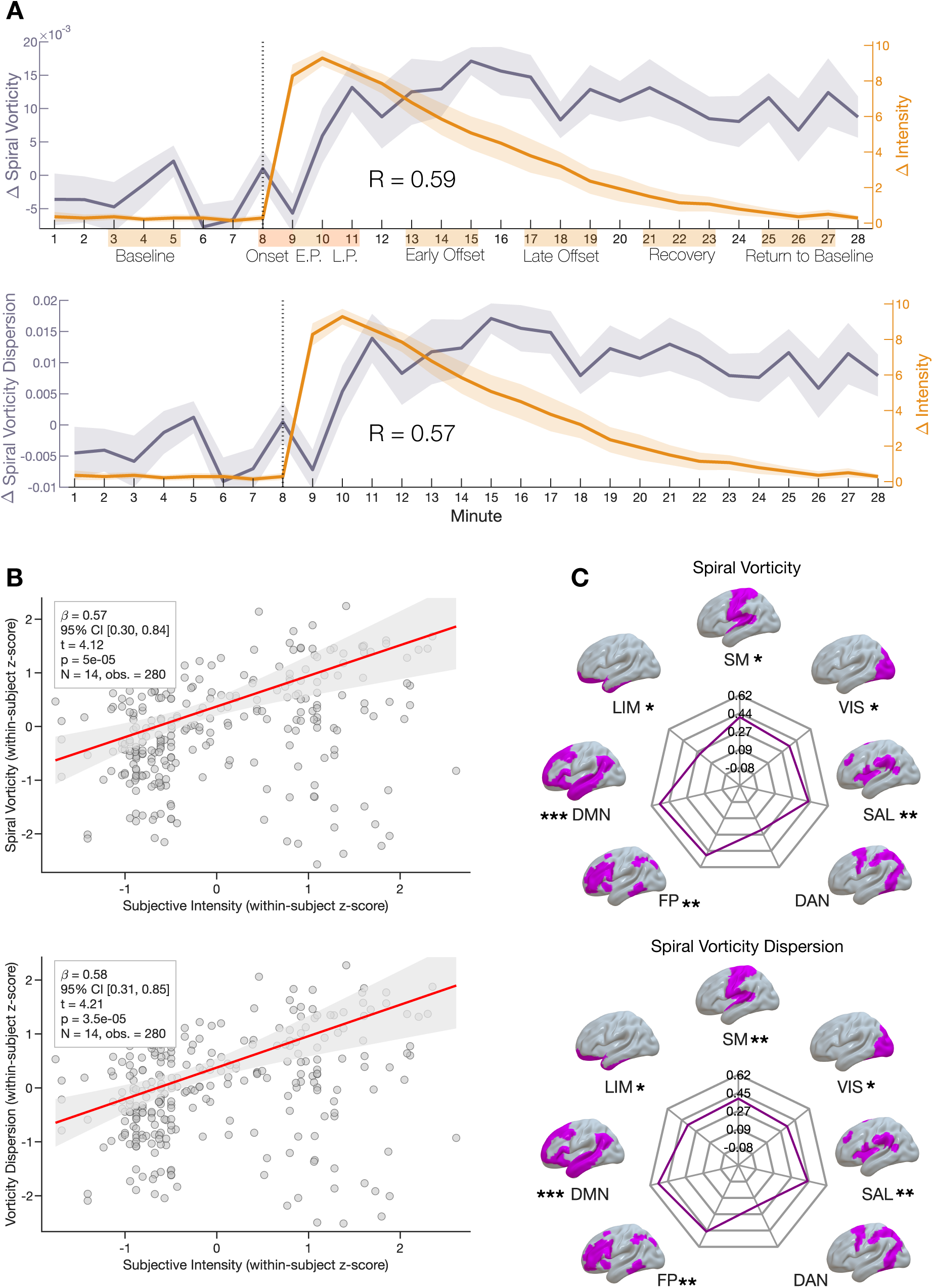
Subjective intensity tracks DMT-induced changes in spiral dynamics. **(A) Global temporal coupling.** Group-average time courses (mean ± SEM) of the DMT–PCB difference in (i) Spiral Vorticity (SV) and (ii) Spatial Spiral Vorticity Dispersion (SSVD), overlaid with the corresponding DMT–PCB difference in subjective intensity ratings (0–10 scale). Subjective intensity was positively correlated with both spiral metrics (SV: Spearman’s *ρ* = 0.59, *p_perm_* = 0.0016; SSVD: *ρ* = 0.57, *p_perm_* = 0.0019; *n* = 28 time points; both BH FDR-corrected across the two metrics, *q* = 0.05 ). The x-axis indicates the experiential phases used in the subsequent spatiotemporal analyses (Figure 4). **(B) Global brain– experience coupling.** Linear mixed-effects models relating within-subject fluctuations in subjective intensity to global neural dynamics during the post-DMT period (minutes 9–28). Points represent participant-by-minute observations. Red lines show the fixed-effect relationship after adjusting for linear and quadratic time effects. **(C) Network-level brain–experience coupling.** Linear mixed-effects models relating within-subject fluctuations in subjective intensity to neural dynamics within each Yeo 7 resting-state network during the post-DMT period (minutes 9–28). Asterisks denote BH FDR-corrected significant associations across the seven networks (* *p_FDR_* < 0.05, ** *p_FDR_* < 0.01, *** *p_FDR_* < 0.001). Full network-specific regression plots are provided in the Supplementary Material.

**Figure 4.**
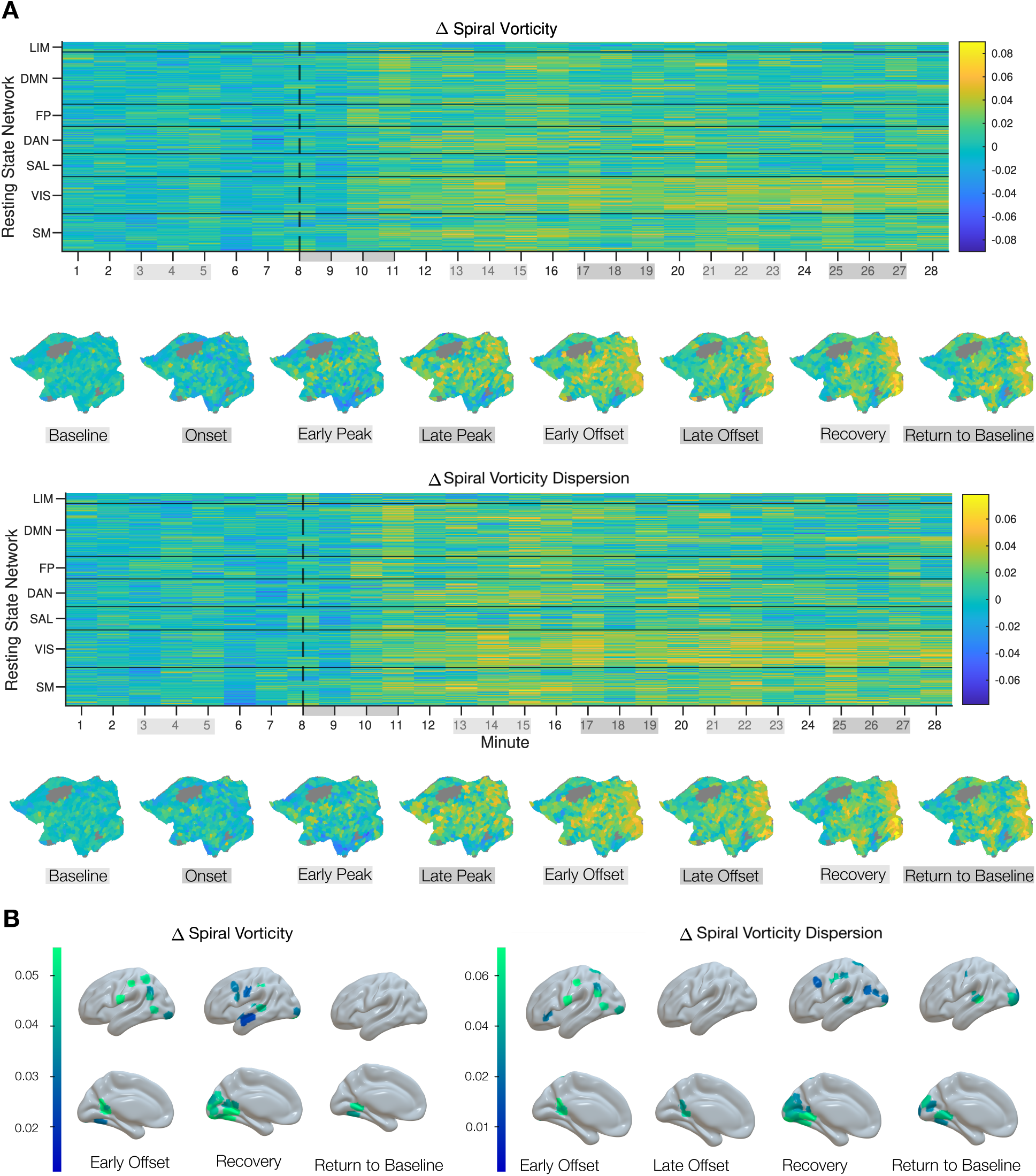
Time-resolved DMT modulation of spiral dynamics across the cortex. **(A) Spatiotemporal evolution of spiral dynamics.** Minute-wise DMT–PCB differences (*Δ*) in Spiral Vorticity (SV; top) and Spatial Spiral Vorticity Dispersion (SSVD; bottom) for each Schaefer1000 parcel, grouped according to the Yeo 7-network parcellation. Heatmaps show parcel-wise changes over time, while stage-averaged cortical flatmaps summarize the spatial distribution of *Δ* values across the eight experiential phases (Baseline, Onset, Early Peak, Late Peak, Early Offset, Late Offset, Recovery, and Return to Baseline). **(B) Stage- and region-specific effects.** Cortical renderings highlight parcels exhibiting significant DMT–PCB differences in Spiral Vorticity (left) and Spatial Spiral Vorticity Dispersion (right) during each experiential stage. Colours represent paired *t*-statistics, with significance determined using paired *t*-tests and Benjamini–Hochberg (BH) FDR correction across the 500 cortical parcels within each stage.

### Experimental paradigm

Participants attended two study days separated by two weeks, each comprising two scanning sessions. The first scan lasted 28 minutes, with intravenous DMT or saline placebo administered at minute 8 as a 60-second bolus. Participants lay in the scanner with their eyes closed and wore an eye mask. Subjective effects were assessed after scanning. The second session followed the same procedure but included minute-by-minute subjective intensity ratings. EEG was recorded simultaneously during both sessions (Figure 1).

### fMRI acquisition parameters

Data were acquired using an EEG-compatible 3T Siemens Magnetom Verio scanner (syngo MR B17). Functional images were collected using a T2^∗^-weighted echo-planar imaging sequence: TR/TE = 2000/30 ms, acquisition time = 28.06 minutes, flip angle = 80°, voxel size = 3 × 3 × 3 mm^3^ , and 35 contiguous slices. High-resolution T1-weighted structural images were also acquired.

### fMRI pre-processing

Preprocessing followed a pipeline developed for a previous LSD study (Carhart-Harris et al. 2016) and included despiking, slice-timing and motion correction, brain extraction, rigid-body registration to structural images, nonlinear registration to a 2-mm MNI template, motion scrubbing, 6-mm FWHM spatial smoothing, 0.01–0.08-Hz band-pass filtering, linear and quadratic detrending, and regression of nine nuisance parameters comprising three translations, three rotations, and three anatomical signals. Preprocessed time series were projected from MNI volumetric space to HCP surface-vertex space using the HCP volume-to-surface procedure.

### Interpolation to a 2D flatmap grid

Vertex-wise fMRI time series were interpolated onto a downsampled 2D cortical flatmap using cubic interpolation, enabling spatial field analyses on a regular grid. A cortical mask derived from the flattened surface geometry was applied to exclude non-cortical regions. The resulting data were represented as 251 × 176 × 240 (or 840) matrices (*y* × *x* × timepoints) and analysed either at the level of the Schaefer1000 parcellation or grouped into Yeo 7 resting-state networks (RSNs).

To account for intersubject variation in spatial sampling, parcels with ≥ 30% missing vertices were excluded. Robustness analyses used group masks retaining regions represented in ≥ 65% or 100% of participants under both conditions (Supplementary Material). Hemispheres were analysed separately, with left-hemisphere results reported in the main text and right-hemisphere results in the Supplementary Material.

### Grid-wise Hilbert phase computation

Instantaneous phase was calculated independently at each cortical grid location using the Hilbert transform (hilbert in MATLAB), yielding a three-dimensional phase field, *φ*(*x*, *y*, *t*).

### Phase-vector field computation

Spatial phase dynamics were characterised by calculating the gradient of the instantaneous phase field, *φ*(*x*, *y*, *t*), across the flatmap, with phase differences corrected for circular wrapping. Gradient vectors were normalised to unit length to obtain a direction-only phase-flow field:

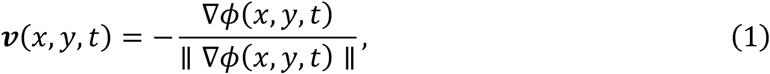

where the negative sign orients vectors in the direction of decreasing phase. This normalised vector field was used to compute the scalar curl, providing a measure of local rotational dynamics at each grid point and timepoint. For visualisation, phase-flow vectors were overlaid on the cortical flatmap as quiver plots (Figure 1C).

### Neural metrics

#### Spiral vorticity

Spiral vorticity (SV) quantifies local rotational flow in the instantaneous cortical phase-flow field. It was calculated as the curl of the normalised phase-vector field:

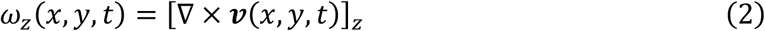

Because ***v*** is defined on a two-dimensional flatmap, only the out-of-plane *z* -component is present. This was computed using MATLAB’s curl function at each time point. Rotation strength irrespective of direction was represented by the absolute vorticity magnitude, |*ω_z_*(*x*, *y*, *t*)|.

### Spatial spiral vorticity dispersion

Spatial spiral vorticity dispersion (SSVD) quantifies the spatial variability of vorticity magnitude. Starting from the absolute vorticity field,

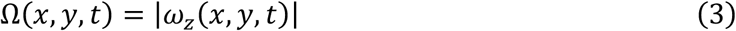

for each cortical parcel *p*, spiral vorticity dispersion was defined as the standard deviation of Ω(*x*, *y*, *t*) across grid locations within that parcel:

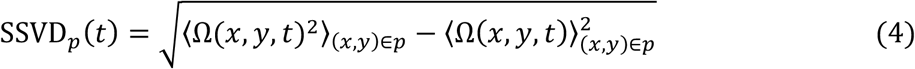

Here, ⟨⋅⟩_(*x*,*y*)∈*p*_denotes averaging across grid locations within parcel *p* at time *t*. Thus, SSVD measures within-parcel spatial dispersion rather than temporal variability.

### Local synchrony and synchrony dispersion

Local phase synchrony was quantified using a neighbourhood Kuramoto order parameter. At each grid location (*x*, *y*) and time *t*, synchrony was calculated as the magnitude of the circular mean of phases within a 3 × 3 neighbourhood:

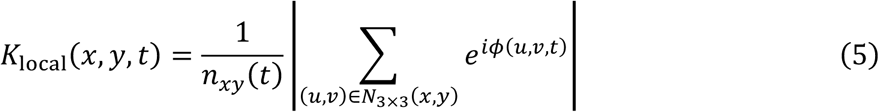

where *n_xy_*(*t*) is the number of valid grid locations in the neighbourhood.

Spatial synchrony dispersion, *K*_SSD_, quantified the spatial variability of local synchrony. For each cortical parcel *p*, it was defined as the standard deviation of *K*_local_(*x*, *y*, *t*) across grid locations within that parcel:

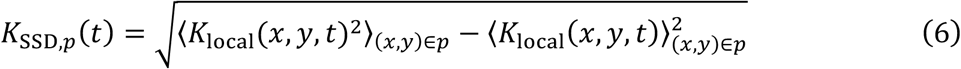

### Temporal irreversibility

Temporal irreversibility in phase dynamics was estimated from the normalised phase-flow field (Equation 1) by calculating directional transition probabilities at each grid location. At location (*i*, *j*) and time *t*, the flow direction was defined as

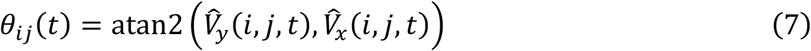

and wrapped to the interval [0,2*π*] for circular binning. Directions were assigned to eight angular bins corresponding to neighbouring grid locations. Valid directional transitions were accumulated over time and normalised to produce a grid-by-eight transition-probability matrix for each participant.

Under detailed balance, the probability of a transition from location *A* to *B* equals that of the reverse transition from *B* to *A*; deviations from this symmetry indicate irreversible dynamics (Figure 1D). Breaking of detailed balance (BDB) at location *A* was therefore calculated as

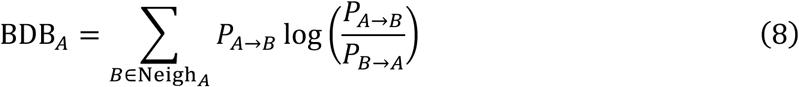

### Statistical analysis

Statistical analyses were performed in MATLAB R2024b (MathWorks, Natick, MA, USA). Global and network-level differences between experimental conditions were assessed using paired sign-flip permutation tests. For parcel-wise analyses, paired *t*-tests were used to compare DMT and placebo conditions at each cortical parcel within each experiential stage. Correlations between subjective intensity and neural dynamics were assessed using Spearman’s rank correlation coefficient, with statistical significance determined using permutation testing. Subject-level correlations were evaluated using two-sided Wilcoxon signed-rank tests.

To account for multiple comparisons, *p*-values were corrected using the Benjamini–Hochberg false discovery rate (BH–FDR) procedure. Unless otherwise stated, significance was defined as *p*_FDR_ < 0.05.

To examine within-subject relationships between subjective intensity and neural dynamics while accounting for the characteristic temporal profile of the DMT response, linear mixed-effects models were fitted to minute-wise observations during the post-infusion period (minutes 9–28). Subjective intensity and neural metrics were *z* -scored within participant, such that model coefficients reflected within-subject deviations from each participant’s post-DMT mean. Fixed effects included subjective intensity together with linear and quadratic time terms, while participant was modelled as a random intercept. Separate models were fitted at the whole-brain and resting-state-network levels.

## Results

We leveraged the predictable time course and robust effects of DMT on conscious experience and brain activity to examine cortical phase dynamics during the acute psychedelic state. We applied our framework to fMRI data acquired during DMT and placebo sessions in a within-subject, placebo-controlled design (**Figure 1A**). BOLD activity was projected onto a two-dimensional cortical flatmap, and the instantaneous phase of each grid element was estimated using the Hilbert transform to generate a time-resolved cortical phase field (**Figure 1B**).

Our primary analyses focused on two vorticity-based measures: spiral vorticity (SV) and spatial spiral vorticity dispersion (SSVD). Spatial phase gradients were used to construct a phase-gradient vector field, from which SV quantified local rotational structure. SSVD quantified the spatial dispersion of cortical vorticity values, indexing heterogeneity in rotational organisation across the cortex (**Figure 1C**). Both measures yielded scalar fields that could be mapped across the cortex and averaged within atlas parcels or resting-state networks.

We also quantified temporal irreversibility (TI) from asymmetries between forward and reverse phase-flow transitions, providing a measure of broken detailed balance in cortical phase dynamics (**Figure 1D**). Because TI was estimated over temporal windows rather than at individual time points, it was not analysed minute by minute.

Our main analyses tested whether DMT-induced changes in brain dynamics tracked subjective intensity (**Figure 1E**). Correlational analyses assessed temporal alignment between SV, SSVD, and intensity (**Figure 1E, left**), whereas network-specific linear mixed-effects models assessed spatial variation in brain–experience coupling (**Figure 1E, right**).

### DMT induces global and network-specific changes in brain dynamics

We quantified SV, SSVD, and TI globally and within the Yeo 7-network parcellation across four conditions: Pre-PCB, Post-PCB, Pre-DMT, and Post-DMT.

Globally, Post-DMT showed robust increases in SV and SSVD relative to all other conditions (**Figure 2A**). SV was higher than in Pre-DMT, Post-PCB, and Pre-PCB ( *p*_FDR_ = 0.0012 , 0.0028, and 0.0012, respectively), as was SSVD (*p*_FDR_ = 0.0030, 0.0108, and 0.0030). By contrast, TI was lower in Post-DMT than in Pre-DMT, Post-PCB, and Pre-PCB (*p*_FDR_ = 0.0036, 0.0064, and 0.0024). Thus, DMT increased the magnitude and spatial dispersion of rotational phase-flow dynamics while reducing temporal irreversibility.

These effects varied across cortical networks (**Figure 2B**). The most consistent SV increases occurred in the default mode network (DMN) and dorsal attention network (DAN), where Post-DMT differed from all other conditions. In the DMN, SV was increased relative to Pre-DMT, Post-PCB, and Pre-PCB (all *p*_FDR_ = 0.0028 ). The DAN showed a similar pattern, with significant Post-DMT increases relative to Pre-DMT, Post-PCB, and Pre-PCB (*p*_FDR_ = 0.0028, 0.0063 , and 0.0028 ). SV was also increased in the visual network (VIS; *p*_FDR_ = 0.0063 , 0.0160, and 0.0065) and frontoparietal network (FPN; *p*_FDR_ = 0.0028, 0.0265, and 0.0092) relative to the same respective conditions. In the somatomotor network (SMN), SV was increased in Post-DMT relative to Pre-DMT and Pre-PCB (*p*_FDR_ = 0.0265 and *p*_FDR_ = 0.0300), but not relative to Post-PCB.

SSVD showed a similar network profile. The strongest and most consistent effects were again observed in the DMN, where Post-DMT was significantly increased relative to all other conditions (all *p*_FDR_ = 0.0042). Increases were also observed relative to Pre-DMT, Post-PCB, and Pre-PCB in the DAN (*p*_FDR_ = 0.0084, 0.0149, and 0.0096) and VIS (*p*_FDR_ = 0.0084, 0.0280, and 0.0126). In the FPN, SSVD was increased in Post-DMT relative to Pre-DMT and Pre-PCB (*p*_FDR_ = 0.0042 and *p*_FDR_ = 0.0280), whereas in the SMN only the comparison with Pre-DMT survived correction (*p*_FDR_ = 0.0280).

Network-level TI effects were more spatially restricted. The strongest decrease was localised to the DMN, where Post-DMT was significantly lower than Pre-DMT, Post-PCB, and Pre-PCB (all *p*_FDR_ = 0.0084). TI was also lower in the FPN relative to Post-PCB (*p*_FDR_ = 0.0315). No FDR-significant effects were observed in the SMN, VIS, salience network (SAL), DAN, or limbic network (LIM) after correction across the 42 resting-state-network-level comparisons per metric.

Together, these results show that DMT globally increases spiral vorticity and its spatial dispersion while reducing temporal irreversibility. Network-level effects were concentrated in higher-order association, attentional, and perceptual systems rather than distributed uniformly across the cortex. Comparable effects were observed in the right hemisphere (**Figure S2**).

### DMT-induced changes in spiral dynamics track subjective intensity

We used complementary correlational and mixed-effects analyses to test whether DMT-related changes in neural dynamics tracked subjective intensity over time.

First, we examined whether drug-specific changes in spiral dynamics followed the temporal profile of subjective intensity. Minute-wise DMT–PCB differences were calculated for subjective intensity and each neural metric and then averaged across participants. Both ΔSV and Δ SSVD were positively associated with Δ intensity over time (**Figure 3A**; SV: *ρ* = 0.59 , *p*_perm_ = 0.0016 ; SSVD: *ρ* = 0.57 , *p*_perm_ = 0.0019 ). Minutes with stronger DMT-related subjective effects therefore showed larger increases in spiral vorticity and its spatial dispersion. Because these analyses used subject-averaged trajectories, they demonstrate group-level temporal alignment rather than within-subject coupling.

Next, we assessed whether within-subject, minute-to-minute fluctuations in subjective intensity covaried with fluctuations in spiral-based neural dynamics across the 28-minute DMT session. Permutation testing indicated significant associations in 6 of 14 participants for SV and 7 of 14 participants for SSVD (Spearman correlations with permutation-based *p* -values, BH–FDR corrected across subjects). At the group level, the distributions of subject-level *ρ* values differed from zero for both metrics (SV: median *ρ* = 0.419, *p* = 0.00045; SSVD: median *ρ* = 0.419, *p* = 0.00045; two-sided Wilcoxon signed-rank tests with BH–FDR correction across metrics). These findings indicate that moment-to-moment subjective intensity covaries with both vorticity-based measures.

To determine whether this coupling remained after accounting for the characteristic temporal trajectory of the DMT response, we fitted linear mixed-effects models to the post-infusion window (minutes 9–28). Models were fitted to participant-by-minute observations using within-subject *z*-scored subjective intensity and neural metrics, such that values reflected deviations from each participant’s own post-DMT mean. Linear and quadratic post-injection time terms were included as covariates, with subject included as a random intercept (280 observations; 14 participants).

Globally (**Figure 3B**), within-subject subjective intensity was positively associated with both SV (*β* = 0.571, SE = 0.139, *t*(275) = 4.12, *p* = 5.00 × 10^−5^ ; 95% CI [0.30,0.84]) and SSVD (*β* = 0.582, SE = 0.138, *t*(275) = 4.21, *p* = 3.51 × 10^−5^; 95% CI [0.31,0.85]). Thus, within-participant increases in subjective intensity were associated with stronger spiral dynamics beyond the average linear and nonlinear temporal profile of the DMT response.

Finally, we repeated the time-controlled mixed-effects analysis within each Yeo 7 resting-state network (**Figure 3C**; full regression plots are provided in the Supplementary Material). For each network, we tested whether within-subject fluctuations in subjective intensity predicted within-subject fluctuations in neural dynamics during the post-DMT period. Intensity positively predicted SV in the DMN (*p*_FDR_ < 0.001), FPN (*p*_FDR_ = 0.002), SAL (*p*_FDR_ = 0.007), SMN (*p*_FDR_ = 0.013), LIM (*p*_FDR_ = 0.017), and VIS (*p*_FDR_ = 0.020), but not in the DAN (*p*_FDR_ = 0.214). A similar pattern was observed for SSVD, with significant associations in the DMN ( *p*_FDR_ = 0.001 ), FPN ( *p*_FDR_ = 0.002 ), SMN ( *p*_FDR_ = 0.008 ), SAL ( *p*_FDR_ = 0.008 ), LIM (*p*_FDR_ = 0.027), and VIS (*p*_FDR_ = 0.028), but not in the DAN (*p*_FDR_ = 0.165).

Brain–experience coupling was therefore widespread but heterogeneous, with the strongest effects consistently observed in the DMN and FPN. Notably, although the DAN showed condition-level DMT effects (**Figure 2B**), its dynamics did not covary significantly with moment-to-moment intensity. Thus, not all attention-related systems tracked the subjective intensity trajectory similarly.

### Time-resolved DMT modulation of spiral dynamics across the cortex

Spatiotemporal heatmaps of minute-wise DMT–PCB differences showed time-dependent modulation of spiral dynamics across cortical parcels (**Figure 4A**). During the pre-infusion baseline period, differences in both SV and SSVD were relatively small and centred near zero. Following infusion, both shifted towards positive DMT–PCB differences across multiple networks, with the most consistent increases observed in visual-network parcels. Stage-averaged cortical flatmaps showed that these effects became more spatially extensive after the late-peak phase, persisted into recovery, and became more restricted during the return-to-baseline period.

Parcel-wise statistics confirmed that these effects were stage- and region-dependent (**Figure 4B**). Significant SV effects occurred during early offset, recovery, and return to baseline, whereas SSVD effects also extended into late offset. During early offset, both metrics differed significantly across visual and default-mode regions, with additional dorsal-attention and somatomotor involvement (VIS: SV = 5 parcels, SSVD = 3 parcels; DMN: SV = 4, SSVD = 5; DAN: SV = 2, SSVD = 2; SMN: SV = 2, SSVD = 2; all FDR-corrected *p* < 0.05).

During recovery, effects were dominated by visual parcels. Significant SV effects occurred in VIS (13 parcels), with additional effects in the DMN (4), SMN (3), DAN (1), and FPN (1). SSVD effects were concentrated in VIS (17 parcels), with additional involvement of the DAN (4), DMN (2), and SMN (1; all FDR-corrected *p* < 0.05). By the return-to-baseline period, SV effects were restricted to two visual parcels, whereas SSVD remained more spatially distributed across VIS (9 parcels), SMN (3), and DMN (1) (all FDR-corrected *p* < 0.05).

Together, these findings indicate that DMT-induced alterations in spiral dynamics are strongest during post-peak periods and most consistently expressed in the visual cortex. The greater spatial persistence of SSVD relative to SV suggests that DMT produces more sustained changes in the spatial heterogeneity of vorticity than in its mean magnitude.

## Discussion

This study used vorticity-based metrics to characterise the spatiotemporal evolution of large-scale brain dynamics during the DMT experience. Moving beyond static or spatially averaged descriptions, we show that spiral dynamics provide a time-resolved neural signature of subjective intensity, capturing its progression across cortical space and time. DMT induced widespread but spatially heterogeneous reconfigurations of brain dynamics that closely tracked moment-to-moment fluctuations in experience. Parcel-level analyses combined with temporally resolved experiential-stage binning further showed that DMT-related changes in spiral dynamics evolved in a temporally and spatially specific manner rather than being uniformly distributed across the cortex.

Most existing metrics quantify the strength or variability of synchronisation but remain insensitive to the spatial organisation of phase relationships. Vorticity-based measures instead capture the local rotational structure of cortical phase flows, providing access to patterns of activity that are not detected by conventional approaches. Spiral waves are a key example of such patterns and have increasingly been reported across a range of cognitive states (Xu et al. 2023; Deco et al. 2026), including the psychedelic state induced by 5-MeO-DMT (Blackburne et al. 2025).

Here, we show that DMT increases both the magnitude of rotational cortical phase flows, indexed by spiral vorticity, and their spatial heterogeneity, indexed by spatial spiral vorticity dispersion. These findings align with recent work showing that brain activity under 5-MeO-DMT is organised into structured, large-scale spatiotemporal wave patterns rather than reflecting unstructured variability in EEG signals (Blackburne et al. 2025). Whereas previous studies have primarily characterised the emergence, propagation, and persistence of such waves, including the identification of phase singularities as organising centres (Blackburne et al. 2025; Xu et al. 2023), our approach directly quantifies the local rotational structure of cortical phase flows and their variability across the cortical surface, moving towards a field-based description of cortical dynamics.

The spatial distribution of these effects is also noteworthy. Increases in spiral vorticity and its dispersion were strongest in the default mode network (DMN) and dorsal attention network (DAN), with additional involvement of visual and frontoparietal systems. This indicates that DMT does not modulate rotational dynamics uniformly but preferentially engages both transmodal association cortex and perceptual networks. This pattern is consistent with previous work showing disruption of DMN integrity (Gattuso et al. 2022; Palhano-Fontes et al. 2015; Carhart-Harris et al. 2016), altered visual-system dynamics (Alamia et al. 2020), and flattening of hierarchical brain organisation under DMT (Vohryzek, Cabral, et al. 2024).

Interpreted in relation to the metrics, these findings suggest a shift towards stronger yet less spatially uniform dynamical organisation. Spiral vorticity indexes the strength of local rotational phase flows, whereas vorticity dispersion captures their spatial variability. Their joint increase, most prominently in the DMN, therefore points to more rotationally structured yet spatially heterogeneous dynamics rather than a simple increase in entropy. The additional involvement of perceptual and control networks suggests a widespread redistribution of dynamical stability across the cortical hierarchy, consistent with REBUS-like accounts of relaxed top-down constraints (Carhart-Harris and Friston 2019).

In addition to changes in spiral dynamics, DMT was associated with reduced temporal irreversibility. Temporal irreversibility indexes the degree to which brain activity exhibits a preferred direction in time and therefore reflects departures from equilibrium dynamics (Kringelbach et al. 2024). The observed decrease in irreversibility, alongside increases in spiral vorticity and its dispersion, suggests a shift towards more flexible and less temporally constrained dynamical regimes. Together, these findings indicate that DMT reconfigures brain dynamics towards rotationally structured and spatially heterogeneous activity that is less constrained by directional temporal organisation.

A central finding of this study is that DMT-induced changes in spiral dynamics closely tracked the temporal unfolding of subjective intensity. Across complementary analyses, spiral vorticity and dispersion covaried with moment-to-moment fluctuations in intensity and reliably tracked its evolution over time. This coupling is consistent with previous work showing that time-resolved measures of brain dynamics track the DMT experience. Vohryzek, Luppi, and colleagues (Vohryzek, Luppi, et al. 2025) reported that increases in connectome-harmonic repertoire entropy and shifts in energy-spectrum distribution covary with subjective intensity, indicating an expansion and redistribution of the brain’s dynamical repertoire. Similarly, reductions in control energy under DMT have been linked to higher subjective intensity, suggesting a flattening of constraints on state transitions (Singleton et al. 2025). Complementing these studies, our findings suggest that subjective effects under DMT are associated with reorganisation of the spatiotemporal structure of brain activity. The psychedelic state may therefore be characterised not only by an expanded repertoire and reduced energetic constraints, but also by altered patterns of activity flow across the cortex.

Finally, the temporal profile of these effects revealed that maximal reorganisation of spiral dynamics did not coincide with peak subjective intensity but emerged during offset and recovery. This suggests that resolution of the psychedelic state involves continued large-scale reconfiguration of cortical dynamics, consistent with previous reports of evolving brain states under DMT (Timmermann et al. 2019), ketamine, LSD, and psilocybin (Schartner et al. 2017). Together with the parcel-level results, these findings identify recovery from the DMT state as an active and spatially structured process of dynamical re-stabilisation rather than a simple reversal of peak effects. This temporally extended and spatially heterogeneous reorganisation is consistent with previous evidence that psychedelic brain dynamics are both time-dependent and network-specific, with ongoing reconfiguration across the course of the experience (Timmermann et al. 2019; Carhart-Harris et al. 2016; Cruzat et al. 2022).

A limitation of this study is its modest sample size, which may constrain generalisability. However, the consistency of effects across complementary analyses supports their robustness. Replication in larger cohorts will be necessary to establish the stability of these findings and characterise inter-individual variability. More broadly, our results highlight the value of spatiotemporally resolved analyses for capturing dynamic properties of brain activity that are inaccessible to static approaches. Future work should determine whether spiral dynamics reflect specific circuit-level mechanisms or more general properties of large-scale cortical organisation. Extending these methods to modalities with higher temporal resolution, such as MEG or intracranial recordings, and to other altered or pathological states may help establish the generality of these signatures. Spirality-based measures may also provide a potential means of tracking subjective intensity and informing adaptive or personalised dosing strategies in psychedelic research.

## Data Availability Statement

Data can be made available upon request.

## Code Availability

Code underlying the results in this paper is available at: https://github.com/GabaSawi/Spiral_Dynamics_DMT

## Acknowledgements

We thank Elvira Garcia Guzman, Pedro Mediano, and George Blackburne for helpful discussions and valuable suggestions that contributed to the development of the analyses.

## Author Contributions

G.S., M.M.M., J.V., and Y.S.P. conceived and designed the study. R.L.C-H. and C.T. acquired and preprocessed the data. G.S. and M.M.M. performed the analyses and interpreted the results, with input from all authors. J.V., Y.S.P., and G.D. developed the analytical methods and statistical models. G.S. drafted the manuscript, and all authors reviewed and critically revised it.

M.L.K. supervised the study. All authors approved the final version of the manuscript.

## References

Alamia, Andrea, Christopher Timmermann, David J Nutt, Rufin VanRullen, and Robin L Carhart-Harris. 2020. DMT Alters Cortical Travelling Waves. 9 (October): e59784. 10.7554/eLife.59784.

Blackburne, George, Rosalind G. McAlpine, Marco Fabus, et al. 2025. “Complex Slow Waves in the Human Brain Under 5-MeO-DMT.” Cell Reports 44 (8): 116040. 10.1016/j.celrep.2025.116040.

Cabral, Joana, Gustavo Deco, Evgeny Hugues, and Morten L. Kringelbach. 2014. “Exploring the Network Dynamics Underlying Brain Activity During Rest.” Progress in Neurobiology 114: 102–31. 10.1016/j.pneurobio.2013.12.005.

Cabral, Joana, Morten Kringelbach, and Gustavo Deco. 2017. “Functional Connectivity Dynamically Evolves on Multiple Time-Scales over a Static Structural Connectome: Models and Mechanisms.” NeuroImage 160 (March). 10.1016/j.neuroimage.2017.03.045.

Carhart-Harris, R. L., and Karl J. Friston. 2019. “REBUS and the Anarchic Brain: Toward a Unified Model of the Brain Action of Psychedelics.” Pharmacological Reviews 71: 316–44. https://api.semanticscholar.org/CorpusID:195188224.

Carhart-Harris, Robin L., Suresh Muthukumaraswamy, Leor Roseman, et al. 2016. “Neural Correlates of the LSD Experience Revealed by Multimodal Neuroimaging.” Proceedings of the National Academy of Sciences 113 (17): 4853–58. 10.1073/pnas.1518377113.

Cruzat, Josephine, Yonatan Sanz Perl, Anira Escrichs, et al. 2022. “Effects of Classic Psychedelic Drugs on Turbulent Signatures in Brain Dynamics.” Network Neuroscience 6 (4): 1104–24. 10.1162/netn_a_00250.

Das, Anup, Erfan Zabeh, Bard Ermentrout, and Joshua Jacobs. 2026. “Planar, Spiral, and Concentric Traveling Waves Distinguish Behavioral States in Human Memory.” Nature Communications 17: 3542. 10.1038/s41467-026-71386-z.

Deco, Gustavo, and Morten L. Kringelbach. 2020. “Turbulent-Like Dynamics in the Human Brain.” Cell Reports 33 (10): 108471. 10.1016/j.celrep.2020.108471.

Deco, Gustavo, Morten L. Kringelbach, Viktor K. Jirsa, and Petra Ritter. 2017. “The Dynamics of Resting Fluctuations in the Brain: Metastability and Its Dynamical Cortical Core.” Scientific Reports 7 (1): 3095. 10.1038/s41598-017-03073-5.

Deco, Gustavo, Yonatan Sanz Perl, Jianfeng Feng, and Morten L. Kringelbach. 2026. “The Turbulent Brain: Modeling Vortex Interactions for Understanding Human Cognition.” *Network Neuroscience*, March, 1–25. 10.1162/NETN.a.554.

Deco, Gustavo, Yonatan Perl, Samuel Johnson, Niamh Bourke, Robin Carhart-Harris, and Morten Kringelbach. 2024. “Different Hierarchical Reconfigurations in the Brain by Psilocybin and Escitalopram for Depression.” Nature Mental Health 2 (August). 10.1038/s44220-024-00298-y.

Escrichs, Alba, Yamil S. Perl, Camila Uribe, et al. 2022. “Unifying Turbulent Dynamics Framework Distinguishes Different Brain States.” Communications Biology 5 (1): 638. 10.1038/s42003-022-03576-6.

Freeman, Walter J., and James M. Barrie. 2000. “Analysis of Spatial Patterns of Phase in Neocortical Gamma EEGs in Rabbit.” Journal of Neurophysiology 84 (3): 1266–78. 10.1152/jn.2000.84.3.1266.

Garcia-Romeu, Albert, Brennan Kersgaard, and Peter Addy. 2016. “Clinical Applications of Hallucinogens: A Review.” Experimental and Clinical Psychopharmacology 24 (August): 229–68. 10.1037/pha0000084.

Gattuso, James, Daniel Perkins, Simon Ruffell, et al. 2022. “Default Mode Network Modulation by Psychedelics: A Systematic Review.” International Journal of Neuropsychopharmacology 26 (October): 155–88. 10.1093/ijnp/pyac074.

Girn, Manesh, Manoj Doss, Leor Roseman, et al. 2026. “An International Mega-Analysis of Psychedelic Drug Effects on Brain Circuit Function.” Nature Medicine 32 (April). 10.1038/s41591-026-04287-9.

Huang, Xiaoying, Wen Xu, Jian-young Liang, Kentaroh Takagaki, Xin Gao, and Jian-Young Wu. 2010. “Spiral Wave Dynamics in Neocortex.” Neuron 68 (5): 978–90. 10.1016/j.neuron.2010.11.007.

Kringelbach, Morten L., Yonatan Sanz Perl, and Gustavo Deco. 2024. “The Thermodynamics of Mind.” Trends in Cognitive Sciences 28 (6): 568–81. 10.1016/j.tics.2024.03.009.

Lewis-Healey, Evan, Carla Pallavicini, Federico Cavanna, et al. 2026. “Time-Resolved Neural and Experience Dynamics of Medium- and High-Dose n,n-Dimethyltryptamine (DMT).” Journal of Cognitive Neuroscience 38 (June): 1244–63. 10.1162/JOCN.a.2423.

Muller, Lyle, Giovanni Piantoni, Dominik Koller, Sydney S. Cash, Eric Halgren, and Terrence J. Sejnowski. 2016. “Rotating Waves During Human Sleep Spindles Organize Global Patterns of Activity That Repeat Precisely Through the Night.” eLife 5: e17267. 10.7554/eLife.17267.

Palhano-Fontes, Fernanda, Kátia Andrade, Luís Fernando Tófoli, et al. 2015. “The Psychedelic State Induced by Ayahuasca Modulates the Activity and Connectivity of the Default Mode Network.” PLoS ONE 10 (February): e0118143. 10.1371/journal.pone.0118143.

Pasquini, Lorenzo, Jakub Vohryzek, Anira Escrichs, et al. 2024. Long-Term Effects of Psilocybin on Dynamic and Effectivity Connectivity of Fronto-Striatal-Thalamic Circuits. 10.1101/2024.11.06.622302.

Preti, Maria Giulia, Thomas A. W. Bolton, and Dimitri Van De Ville. 2017. “The Dynamic Functional Connectome: State-of-the-Art and Perspectives.” NeuroImage 160: 41–54. 10.1016/j.neuroimage.2016.12.061.

Ribary, U., A. A. Ioannides, K. D. Singh, et al. 1991. “Magnetic Field Tomography of Coherent Thalamocortical 40-Hz Oscillations in Humans.” Proceedings of the National Academy of Sciences of the United States of America 88 (24): 11037–41. 10.1073/pnas.88.24.11037.

Rubino, Doug, Kay Robbins, and Nicholas Hatsopoulos. 2007. “Propagating Waves Mediate Information Transfer in the Motor Cortex.” Nature Neuroscience 9 (January): 1549–57. 10.1038/nn1802.

Schartner, Michael, Robin Carhart-Harris, Adam Barrett, Anil Seth, and Suresh Muthukumaraswamy. 2017. “Increased Spontaneous MEG Signal Diversity for Psychoactive Doses of Ketamine, LSD and Psilocybin.” Scientific Reports 7 (April): 46421. 10.1038/srep46421.

Singleton, S. Parker, Christopher Timmermann, Andrea Luppi, et al. 2025. “Network Control Energy Reductions Under DMT Relate to Serotonin Receptors, Signal Diversity, and Subjective Experience.” Communications Biology 8 (April). 10.1038/s42003-025-08078-9.

Strassman, Rick J. 1995. “Human Psychopharmacology of n,n-Dimethyltryptamine.” Behavioural Brain Research 73 (1): 121–24. 10.1016/0166-4328(96)00081-2.

Timmermann, Christopher, Leor Roseman, Sharad Haridas, et al. 2023. “Human Brain Effects of DMT Assessed via EEG-fMRI.” Proceedings of the National Academy of Sciences of the United States of America 120 (March): e2218949120. 10.1073/pnas.2218949120.

Timmermann, Christopher, Leor Roseman, Michael Schartner, et al. 2019. “Neural Correlates of the DMT Experience Assessed with Multivariate EEG.” Scientific Reports 9 (November): 16324. 10.1038/s41598-019-51974-4.

Townsend, Pulin, Rory G. AND Gong. 2018. “Detection and Analysis of Spatiotemporal Patterns in Brain Activity.” PLOS Computational Biology 14 (December): 1–29. 10.1371/journal.pcbi.1006643.

Townsend, Rory G., Selina S. Solomon, Spencer C. Chen, et al. 2015. “Emergence of Complex Wave Patterns in Primate Cerebral Cortex.” Journal of Neuroscience 35 (11): 4657–62. 10.1523/JNEUROSCI.4509-14.2015.

Vidaurre, Diego, Stephen M. Smith, and Mark W. Woolrich. 2017. “Brain Network Dynamics Are Hierarchically Organized in Time.” Proceedings of the National Academy of Sciences 114 (48): 12827–32. 10.1073/pnas.1705120114.

Vogt, Severin, Laura Ley, Livio Erne, et al. 2023. “Acute Effects of Intravenous DMT in a Randomized Placebo-Controlled Study in Healthy Participants.” Translational Psychiatry 13 (May). 10.1038/s41398-023-02477-4.

Vohryzek, Jakub, Joana Cabral, Christopher Timmermann, et al. 2024. “The Flattening of Spacetime Hierarchy of the n,n-Dimethyltryptamine Brain State Is Characterized by Harmonic Decomposition of Spacetime (HADES) Framework.” National Science Review 11 (5): nwae124. 10.1093/nsr/nwae124.

Vohryzek, Jakub, Elvira Garcia-Guzmán, Morten L. Kringelbach, et al. 2025. “Brain Dynamics of Classical Psychedelics Show Paradoxical Hierarchical Flattening with Increased Complexity.” bioRxiv, ahead of print. 10.1101/2024.12.21.629922.

Vohryzek, Jakub, Morten Kringelbach, Edmundo Lopez-Sola, et al. 2024. Collapse of Directed Functional Hierarchy Under Classical Serotonergic Psychedelics. 10.1101/2024.12.21.629922.

Vohryzek, Jakub, Andrea Luppi, Selen Atasoy, et al. 2025. “N,n-Dimethyltryptamine Effects on Connectome Harmonics, Subjective Experience and Comparative Psychedelic Experiences.” Neuropsychopharmacology : Official Publication of the American College of Neuropsychopharmacology 50 (September). 10.1038/s41386-025-02190-4.

Wu, Jian-Young, Xiaoying Huang, and Chuan Zhang. 2008. “Propagating Waves of Activity in the Neocortex: What They Are, What They Do.” The Neuroscientist 14 (5): 487–502. 10.1177/1073858408317066.

Xu, Yiben, Xian Long, Jianfeng Feng, and Pulin Gong. 2023. “Interacting Spiral Wave Patterns Underlie Complex Brain Dynamics and Are Related to Cognitive Processing.” Nature Human Behaviour 7 (June): 1–20. 10.1038/s41562-023-01626-5.

Ye, Zhiwen, Alexander E. Ladd, Nancy MacKenzie, et al. 2026. “Brain-Wide Topographic Coordination of Rotating Waves.” Science 392 (6804): eadx1369. 10.1126/science.adx1369.

Zalesky, Andrew, Alex Fornito, Luca Cocchi, Leonardo L. Gollo, and Michael Breakspear. 2014. “Time-Resolved Resting-State Brain Networks.” Proceedings of the National Academy of Sciences 111 (28): 10341–46. 10.1073/pnas.1400181111.

