## Supplementary Material for "Spiral brain dynamics track the temporal unfolding of subjective intensity during the *N,N*-dimethyltryptamine experience"

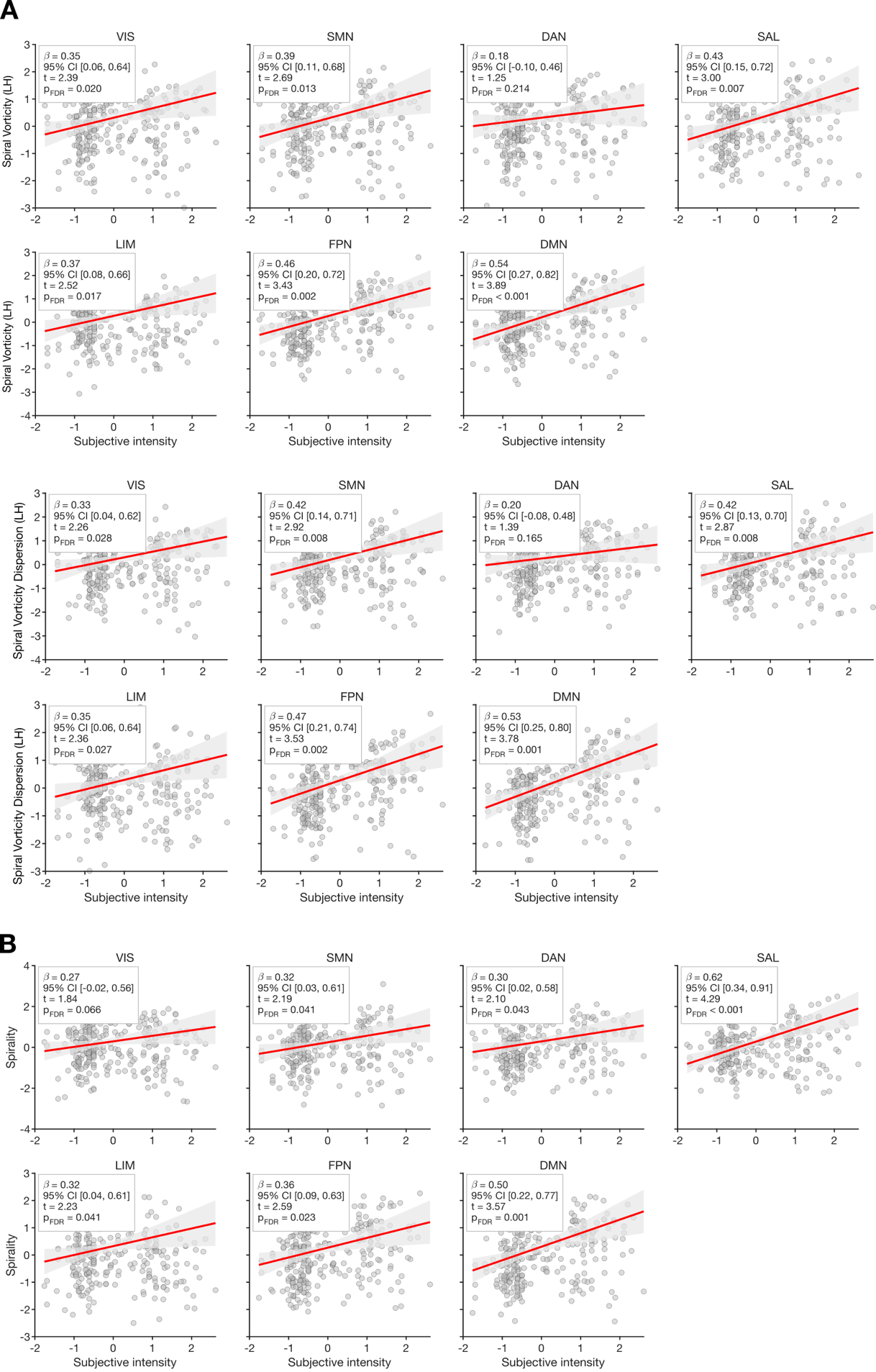
 **Fig. S1** Full linear mixed-effects models regression plots, supplementary to Fig. 3C for each Yeo7 resting-state network (RSN) **(A)** for the left hemisphere (LH) and **(B)** the right hemisphere (RH).

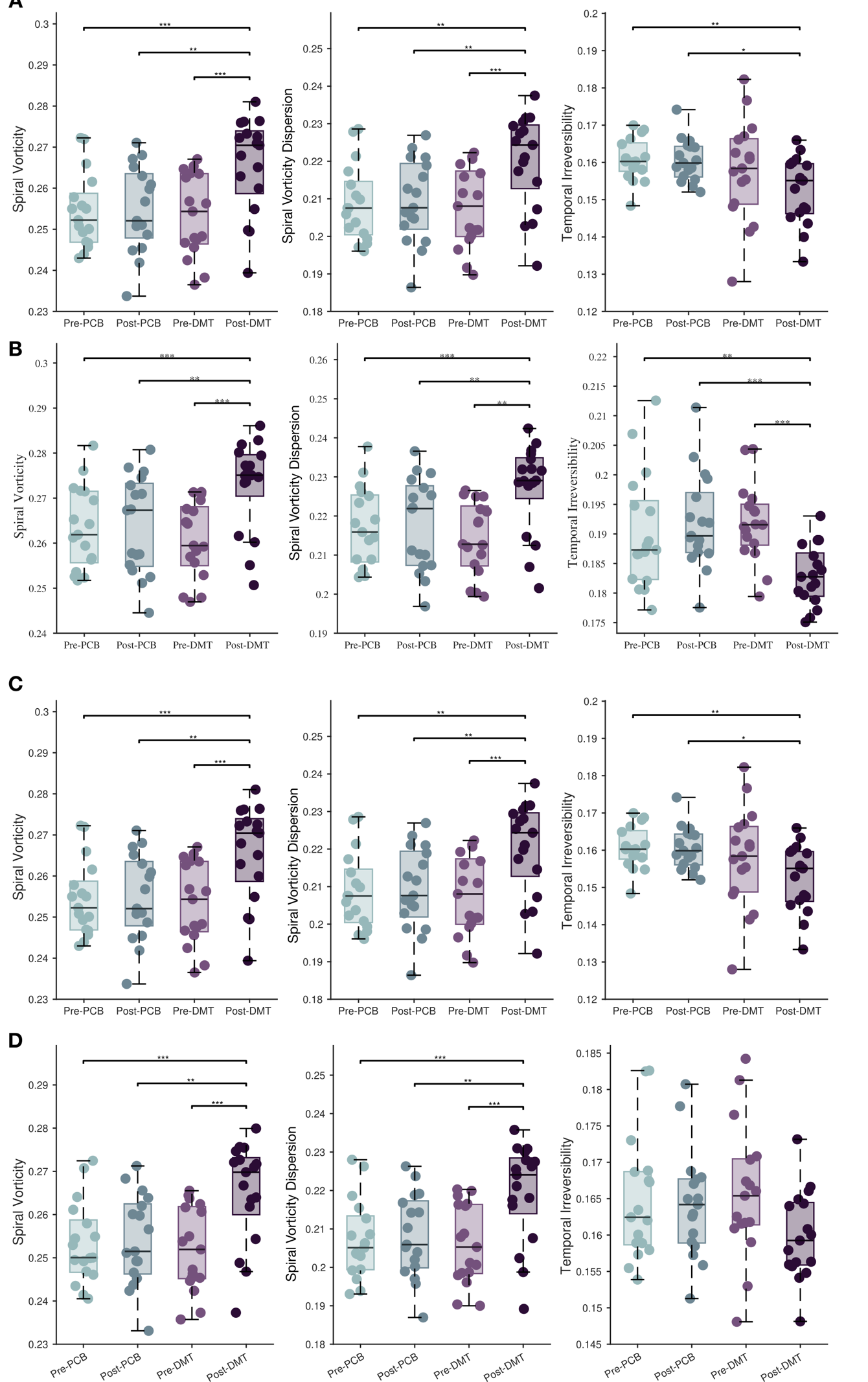

**Fig. S2** Replication of group-level distributions of three neural dynamics metrics under different parcel inclusion thresholds. **(A)** Parcels retained in ≥65% of subjects. **(B)** Parcels retained in 100% of subjects. **(C–D)** Corresponding results for the RH, demonstrating robustness to sample coverage.

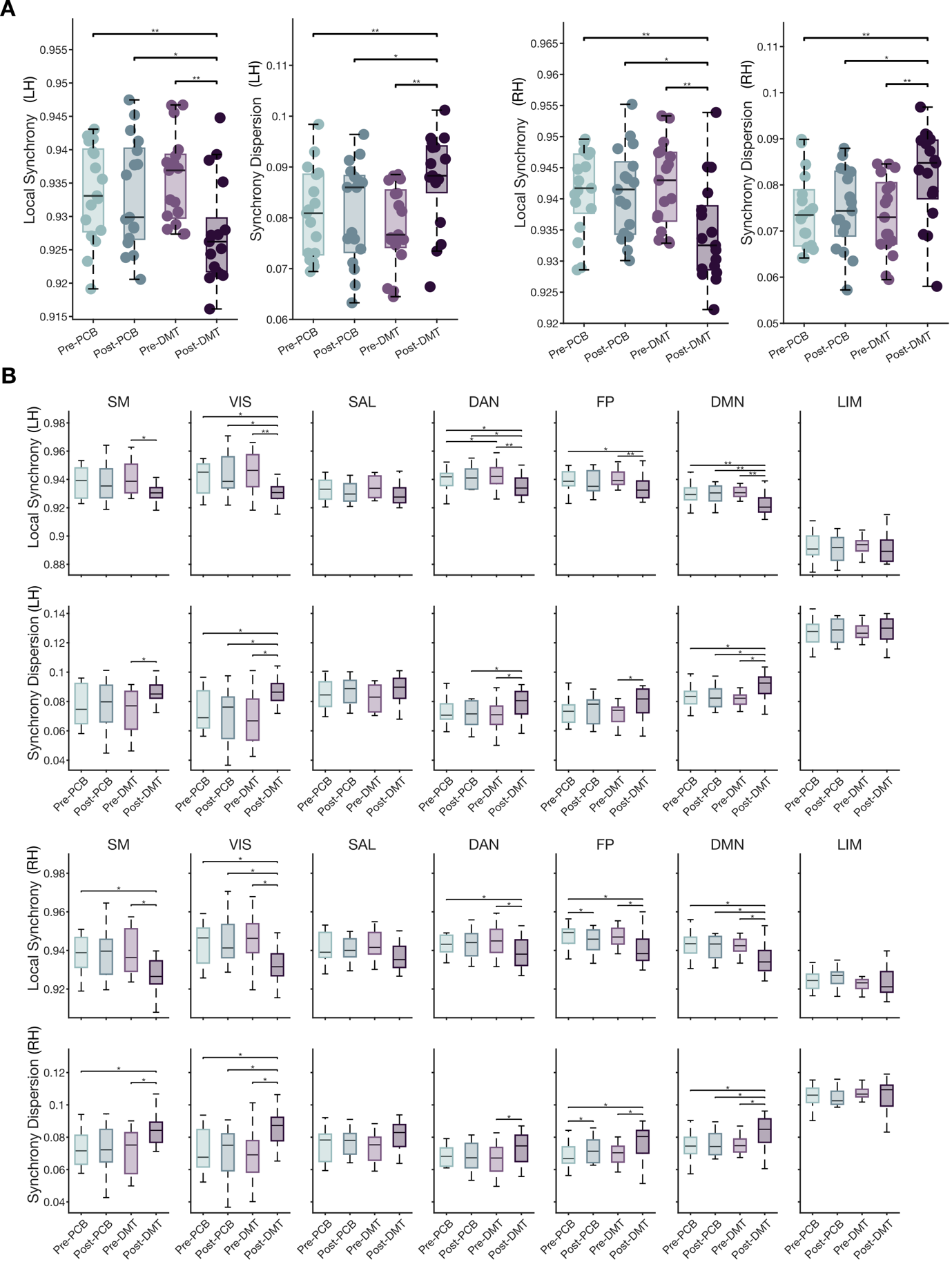

**Fig. S3 (A)** Group-level distributions and **(B)** resting-state network resolved analyses of Kuramoto-based metrics (Kuramoto vorticity and turbulence) for the LH and the RH.

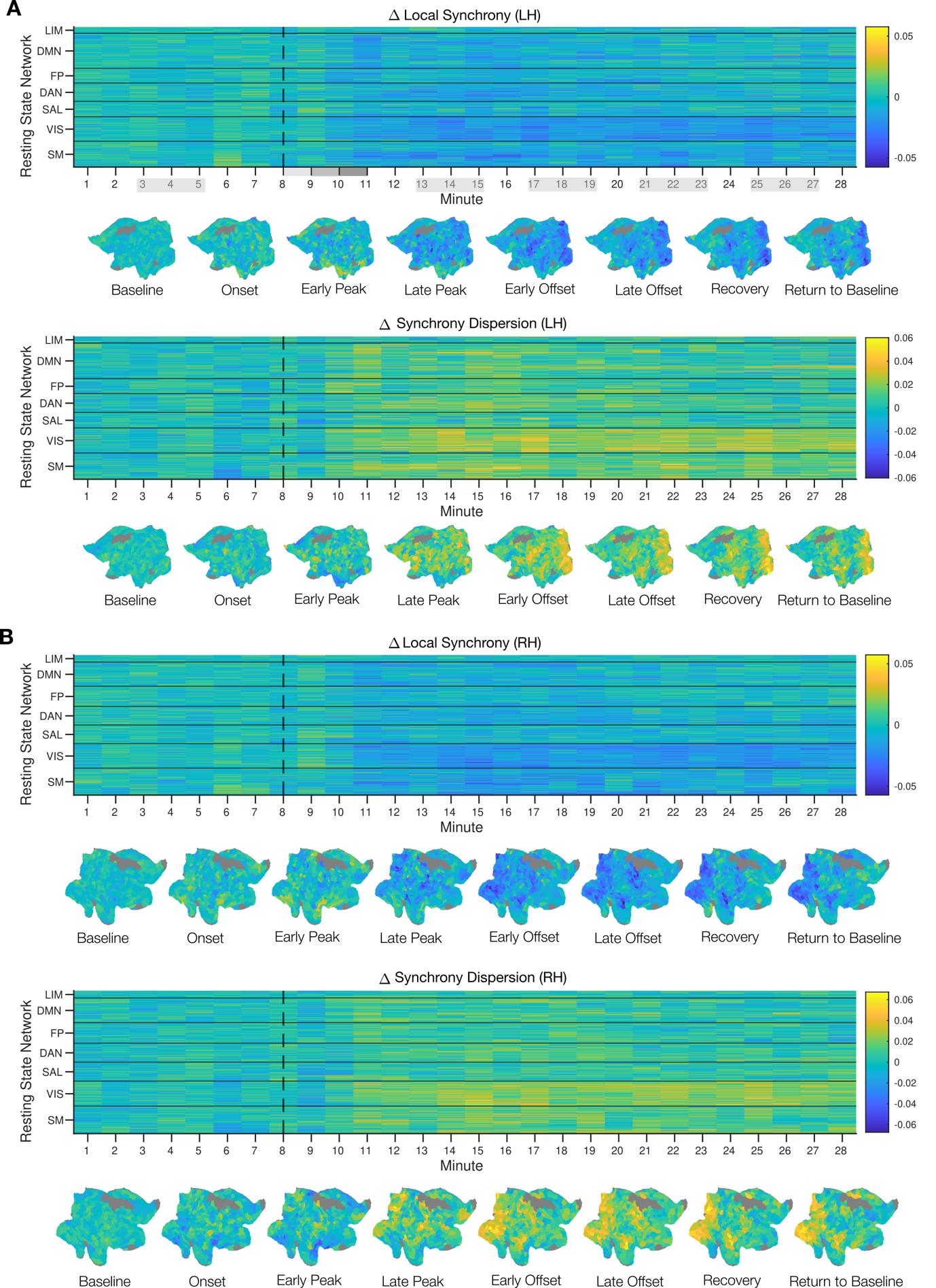

**Fig. S4** Time-resolved DMT modulation of Kuramoto-based dynamics across cortex for **(A)** the LH, and **(B)** the RH.

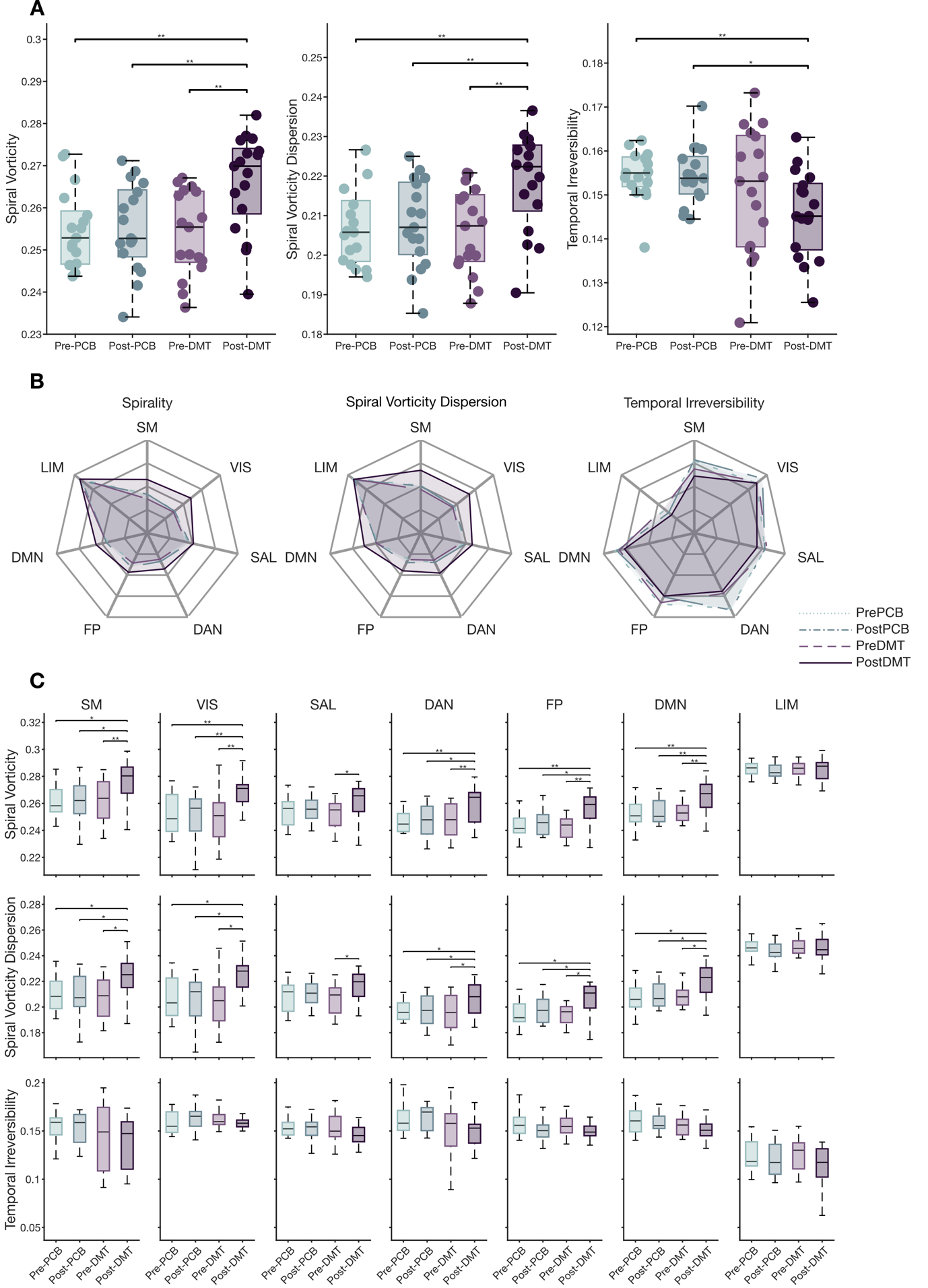

**Fig. S5** Analysis of RH **(A)** group-level distributions and **(B) r**esting-state network resolved analyses of vorticity-based metrics (spiral vorticity and turbulence).

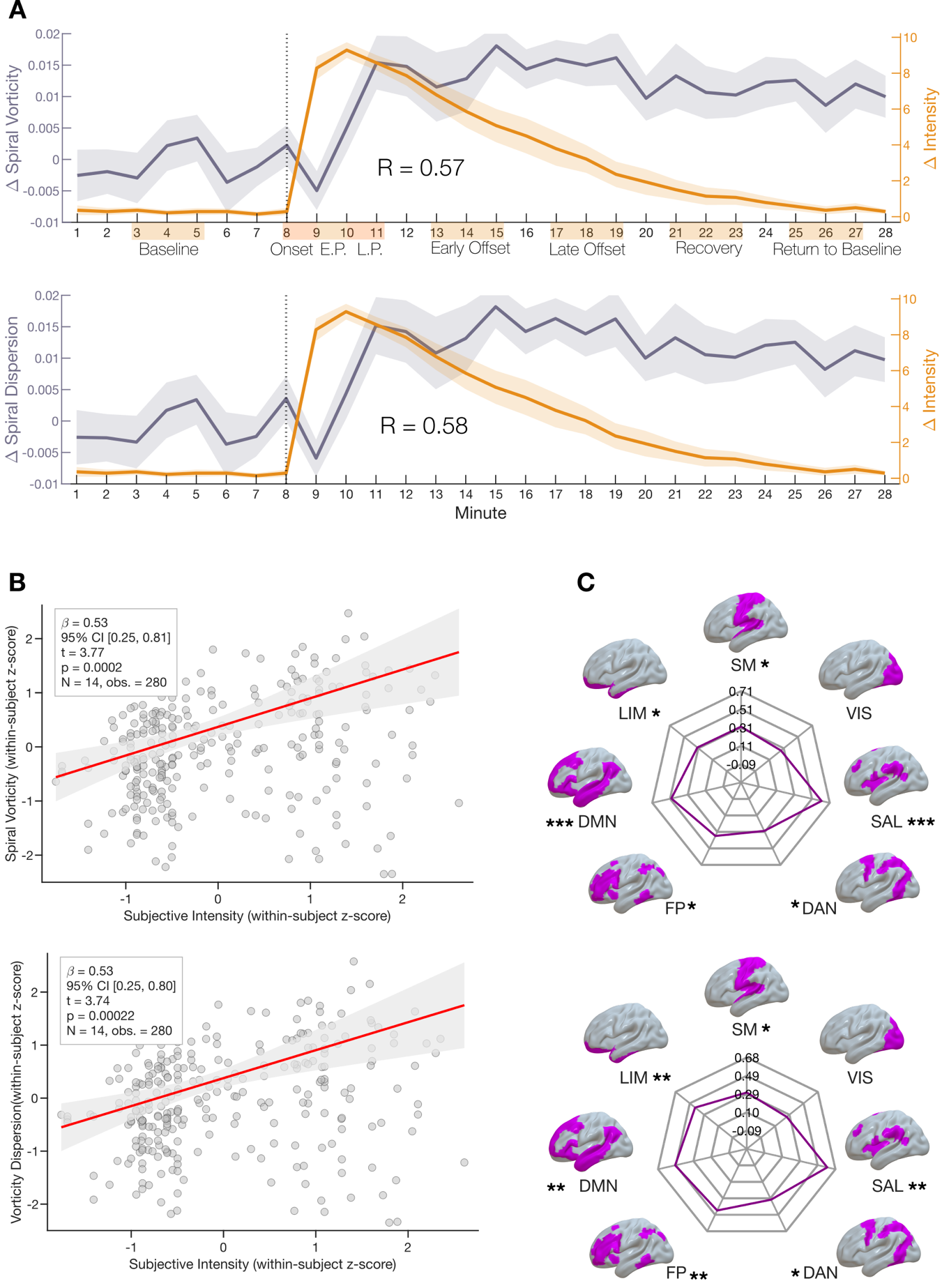

**Fig. S6 Subjective intensity tracks DMT-induced changes in spiral dynamics analysis for the RH (A)** Group-average time courses of the DMT--PCB difference in spiral-based metrics, overlaid with subjective intensity differences (0--10) (N = 28 time points; spiral vorticity: spiral vorticity: R, rho= xx, p = xxx; spiral turbulence: R, rho = xx, p = xxx). **(B)** Mixed-effects relationship between subjective intensity and neural dynamics (xxx). **(C)** Network-level intensity--metric coupling estimated using the same mixed-effects model within each Yeo 7 network. Full RSN-specific models are provided in Fig. S1.

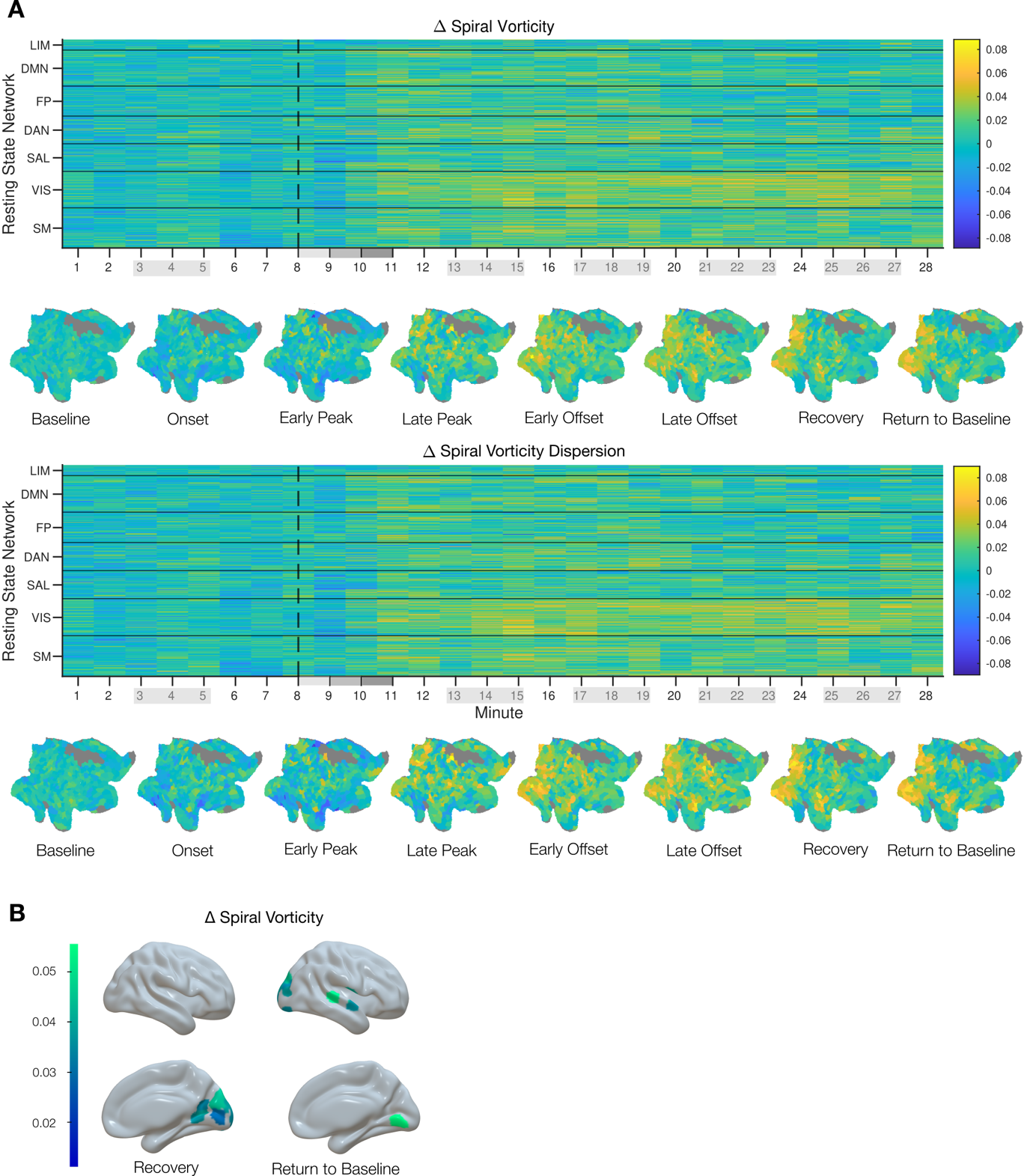

**Fig. S7 RH analysis of time-resolved DMT modulation of spiral dynamics across cortex. (A)** Minute-wise DMT-PCB differences in spiral vorticity (top) and spiral turbulence (bottom) for each Schaefer 1000 parcel and phase-averaged cortical flatmaps summarize changes in patterns across experiential stages. **(B)** Cortical renderings highlighting parcels and stages with significant DMT-PCB differences in spiral vorticity.
